# Advanced median-based genetic similarity analysis in Kazakh Tazy dogs: A novel approach for breed conformity assessment

**DOI:** 10.1101/2024.03.19.585659

**Authors:** Anastassiya Perfilyeva, Rustam Mussabayev, Kira Bespalova, Yelena Kuzovleva, Sergey Bespalov, Mamura Begmanova, Almira Amirgalyeva, Olga Vishnyakova, Inna Nazarenko, Yuliya Perfilyeva, Konstantin Plakhov, Leyla Djansugurova

## Abstract

The breed conformity evaluation is crucial for the preservation of the traits that characterise each dog breed. The use of genetic markers for this purpose provides a precision and objectivity that can surpass the reliability of phenotypic evaluations. In this study, we present a new simple algorithm for assessing breed conformity. The algorithm creates a similarity matrix based on genotypic data and then uses the median to calculate the percentage of genetic similarity that an individual has in relation to the genetic diversity of the breed. To validate the proposed algorithm, we applied it to the genotypic data of 18 microsatellites and 43,691 single nucleotide polymorphisms (SNPs) of the Kazakh Tazy dog, a breed of great cultural and historical importance to Kazakhstan that is now threatened with extinction due to crossbreeding. The algorithm showed a moderate correlation between the microsatellite and SNP genotyping methods, reflecting the different aspects of genetic similarity. In particular, the SNP-based evaluations agreed better with the expert judgements, highlighting their potential for accurate analysis of breed conformity. The proposed algorithm provides easily interpretable results, is flexible, adapts to different genetic markers and may provide an evaluation mechanism for breed conformity in situations where there is no reference population, incomplete pedigrees, unidentified meta-founders and high genetic diversity in the population.

**Author summary:** Our research was initiated by the urgent concern for the possible extinction of the Kazakh Tazy dog, a breed with deep historical roots and cultural significance in Kazakhstan. Information at the DNA level may lead to faster genetic improvement of the breed than relying only on phenotypic data and pedigrees. Genotypic data can be processed to provide valuable insights into genetic diversity, relatedness and ancestry. However, existing methods do not provide a measure of percentage similarity that can be used to assess how closely a particular individual matches the typical genetic composition of the breed. These challenges have led us to propose an approach that overcomes the limitations of existing methods and allows genetic similarity to be assessed based on genotypes. It is based on a median-based approach to analyze genetic data and has been applied to microsatellite and SNP markers but can also be adapted to other genetic markers. This method provides results even in the absence of a defined reference population, complete pedigrees or identified meta-pedigrees and during high genetic diversity within a breed. It can provide breeders and researchers with a tool to maintain the genetic purity of unique breeds.

## Introduction

The Kazakh sighthound dog breed Tazy is not only a unique genetic resource, but also an ancient cultural and historical heritage of Kazakhstan. In 2012, the Law of the Republic of Kazakhstan on the Protection, Reproduction and Utilisation of Wild Animals recognised the status of the Kazakh Tazy breed as a national dog breed. The category of national hunting was also introduced. In 2014, the Ministry of Agriculture of the Republic of Kazakhstan recognised the standard of the Kazakh Tazy breed, which further increased the importance of the breed. Nevertheless, the selection of the breed is still threatened by crossbreeding, which began in the 1950s and intensified after 2014 due to competitions between dog organisations. To improve the running performance, the Kazakh Tazy was crossed with the English Greyhound in the western region, with the Russian Borzoi in the northern region, with the Saluki in the eastern region, with the Saluki and the Kyrgyz Taigan in the southern region and with the Saluki and local dogs in the central region of Kazakhstan. The concept of mandatory bonitation, which was approved by the Ministry of Agriculture in 2014 but has not yet found practical application, could improve the breeding of Kazakh Tazy dogs. It provides for a comprehensive assessment of the dogs based on pedigree, conformation, working ability and quality of offspring. However, traditional evaluations, which focus primarily on phenotypic traits because there are no pedigrees for most Kazakh Tazy dogs, may not capture the full breeding value of the dog because important genetic factors are ignored. This problem can be overcome by using genetic data to improve the reliability of breeding value estimates for dogs in the bonitation process. Information at the DNA level may lead to faster genetic improvement than relying on phenotypic data and pedigrees alone [1–5]

Microsatellites and SNPs are widely used in population genetics and have proven to be suitable markers for analyzing the genetic structure, effective population size and genetic diversity of breeds. Microsatellites are contiguous, highly polymorphic, repetitive DNA segments consisting of two or more nucleotides, while SNPs are genetic markers characterized by variations in the DNA sequence due to changes in a single base in the genome. These markers are commonly used to estimate genetic variation in populations of different dog breeds [6–16]. Researchers have devoted much attention to studying the gene pools of small native dog breeds. The degree of genetic diversity of the Tatra Shepherd Dog [17], the Norwegian Lundehund [9], the Czech Spotted Dog [10]; the Korean Sapsari Dog [18], and the Pungsan and Jindo [19] has been studied. The genetic structure of the Kazakh Tazy has also been characterised using microsatellite and SNP markers [20,21]. These studies emphasise the developing understanding of the role of SNPs and microsatellites in the population genetics of dog breeds. SNPs are increasingly recognised for their frequency, stability, and the possibility of high-throughput genotyping, making them highly valuable for population genetics research. Nevertheless, some studies have shown that microsatellites may be better predictors of genetic relatedness than SNPs [22,23]. Recently, microsatellites were found to show stronger signals than SNPs in discriminating between domestic pigs and wild boars [24]. In addition, a growing number of studies indicate that microsatellites play an important role in controlling evolutionary processes and influence both phenotypic variation and genetic diversity [25]. The choice between SNPs and microsatellites ultimately depends on the specific aims and requirements of the study.

Microsatellite and SNP data can be processed in principal component analysis (PCA), admixture analyses, phylogenetic and other methods to provide valuable insights into genetic diversity, relatedness and ancestry. However, they do not provide a measure of percentage similarity to assess how closely a particular individual matches the typical genetic composition of the breed. These challenges motivate us to propose an approach that overcomes the limitations of existing methods and enables genetic similarity assessment based on genotypes. In this paper, we present the details of the approach used, including algorithmic aspects and implementation considerations. We practically test the algorithm using a microsatellite and SNP dataset of Kazakh Tazy dogs. We also provide an overview of existing methods for calculating genomic breeding values. We discuss the advantages and limitations of the different methods and highlight the gaps in current knowledge that justify the need for the proposed approach.

## Material and methods

### Data

In the study, we used previously published microsatellite genotyping data for DNA samples from 114 Kazakh Tazy dogs [20] as well as additional DNA samples from 109 Kazakh Tazy dowgs collected between May 2022 and May 2023. A detailed description of the biomaterial collection procedure, DNA extraction techniques and microsatellite analysis can be found in our previous study [20].

The dataset used for this study also includes previously published genotyping data for 172,115 SNP markers of the 39 Kazakh Tazy dogs, which represent a subset of a larger cohort of 223 dogs [20]. Plink [26] was used to detect and remove SNPs with more than 10% of missing data (--mind 0.1) and linkage disequilibrium (--indep-pairwise 50 5 0.2), leaving 43,691 SNP markers.

The microsatellite and SNP dataset used in this study can be found in S3 and S4 Tables, respectively.

The study was approved by the Ethics Committee of the Institute of Human and Animal Physiology, Almaty, Kazakhstan (number 3, 15 September 2020).

### Analysis of the microsatellite data

The allele frequencies of 18 microsatellite loci were analysed to calculate the parameters of genetic diversity, such as the average number of alleles per locus (Na), the average number of effective alleles per locus (Ne), the observed heterozygosity (Ho), the expected heterozygosity (He) and the inbreeding coefficient (F) using the GenAIEX 6 software. To analyse the genetic structure of the studied population, a PCA was performed with the software GenAIEX 6.5 and a Bayesian cluster analysis with the software Structure v2.3.4. The following analysis parameters were defined for the Bayesian cluster analysis: Admixture model algorithm, correlation with allele frequency, 10,000 burn-in iterations and 100,000 MCMC repetitions. For each K between 2 and 10, 5 independent analyses were performed. The most probable number of clusters was calculated with the R package Pophelper [27] according to the method of Evanno et al [28].

### Computation of Correspondence Percentage Using Microsatellite Data

Based on the microsatellite genotypes obtained, we developed an algorithm for evaluating the genetic similarity of a particular dog within the Kazakh dog breed Tazy. Its pseudocode is presented in Algorithm 1.

The algorithm operates on a dataset represented as a matrix **G**, where each row ***G***_***i***_ corresponds to the genotypic data of an individual dog. The feature vector **F** of the specific dog under consideration is also provided as input. The pseudocode for the algorithm is as follows:

1. The algorithm initiates by constructing an *F* × *F* similarity matrix **S**, where *N* is the number of individuals in the dataset. This matrix is used to store the genetic similarity scores between pairs of individuals in the population.
2. For each pair of individuals (*i*, *j*), the algorithm calculates the genetic similarity score *S*_*ij*_. This calculation is based on either the Proportion of Shared Alleles (PSA) method or the Jaccard Similarity Index, depending on the method chosen. PSA method involved calculating the proportion of shared alleles at each locus for pairs of individuals. For individuals A and B, the PSA similarity, PSA(A, B), is given by:

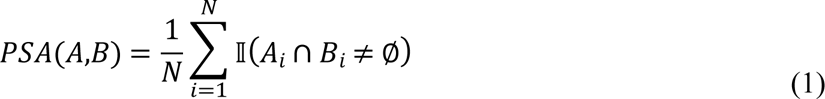 Here, *N* is the total number of loci, *A*_*i*_ and *B*_*i*_ represent the allele sets at the *i*-th locus, and *I* is the indicator function. Jaccard Similarity Index quantifies similarity as the ratio of the intersection to the union of two allele sets. For individuals A and B, the Jaccard similarity, J(A, B), is:

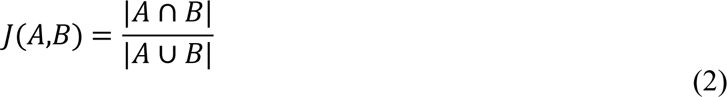
3. The median genetic similarity score *M*_*i*_for each individual is computed by taking the median of the *i*-th row of the similarity matrix **S**.
4. The algorithm then determines the maximum median similarity score *M*_max_ across all individuals, which serves as a reference for typical genetic similarity within the breed:

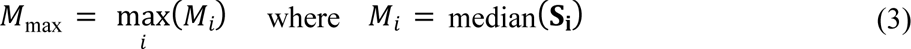
5. A similarity row vector **R** is constructed for the specific individual dog under study, comparing its genotypic data **F** with that of every other individual in the dataset.
6. The median of the similarity row vector **R**, denoted as *M*_ind_, is calculated, representing the median genetic similarity of the specific individual to the rest of the population.
7. Finally, the correspondence percentage *C%* is computed, which quantifies the genetic conformity of the specific individual relative to the typical genetic makeup of the breed. This is achieved by taking the minimum of the ratio of *M*_ind_ to *M*_max_ (capped at 1.0) and multiplying by 100 to express it as a percentage:

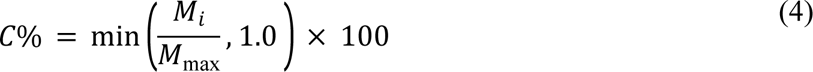

This percentage reflects how an individual’s genetic profile compares to the most common genetic profile in the Tazy breed, serving as a key metric for breed conformity assessment.

#### Algorithm 1 - Correspondence percentage calculation using microsatellite data

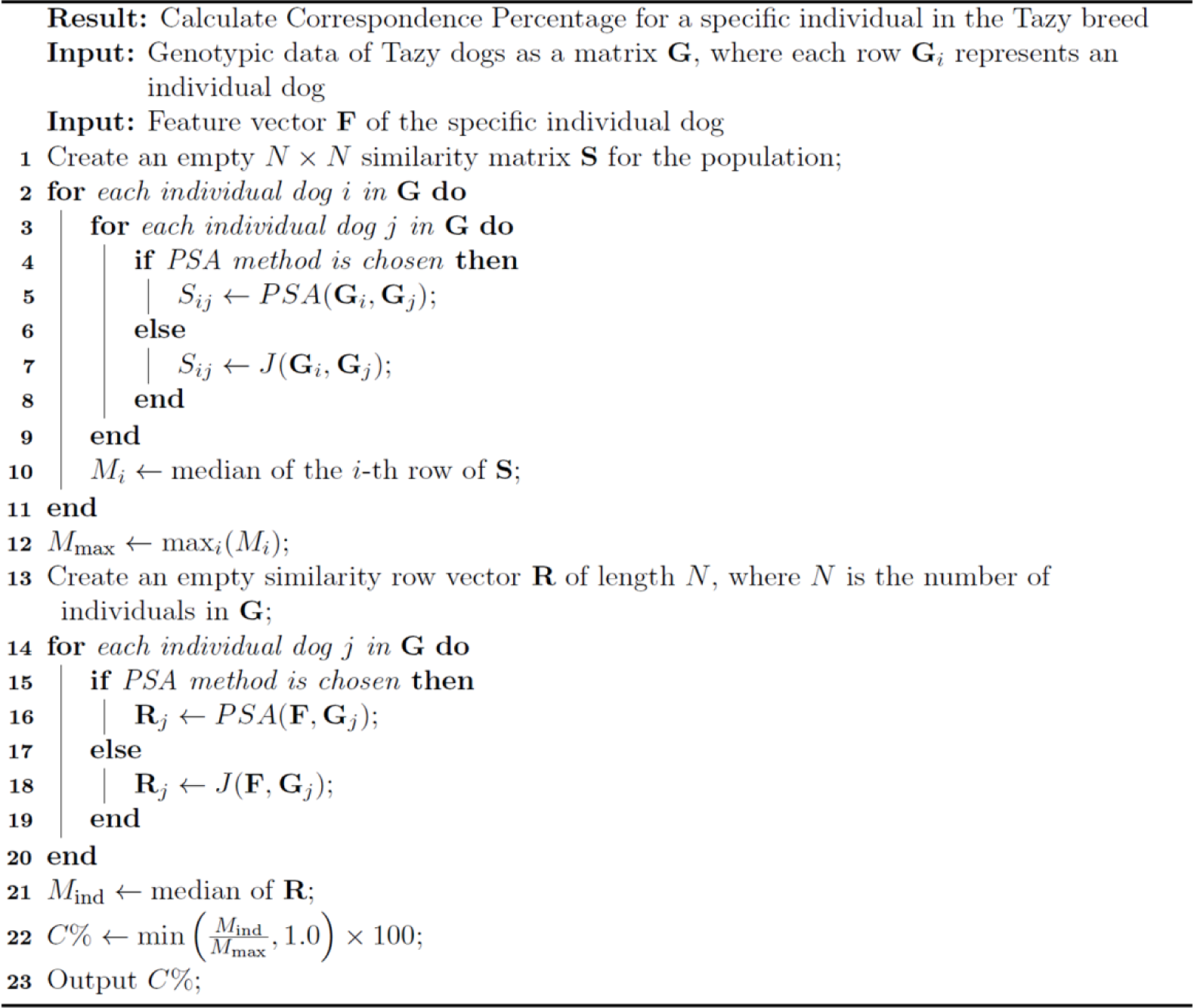

### Computation of Correspondence Percentage Using SNP Data

The proposed algorithm using the multi-allelic microsatellite data was adapted to the bi-allelic genome-wide SNP microarray data of the 39 Kazakh Tazy dogs.

The SNP data for each dog was first subjected to one-hot coding, in which the categorical genotypic information was converted into a numerical format. The possible genotypes, including a ‘missing’ category for absent data, were encoded as follows:

– ‘0/0’ (Homozygous for reference allele) - [1, 0, 0];
– ‘0/1’ (Heterozygous) - [0, 1, 0];
– ‘missing’ - [0, 0, 1].

This process facilitated the calculation of similarity measures and subsequent analysis. The data was then vectorized, with each dog represented as a unique trait vector encompassing its entire genotypic profile. The percentage similarity calculation procedure was then applied using pairwise cosine similarity based on SNP data. The pseudo code of the above process is in Algorithm 2.

#### Algorithm 2 - Correspondence percentage calculation using SNP data

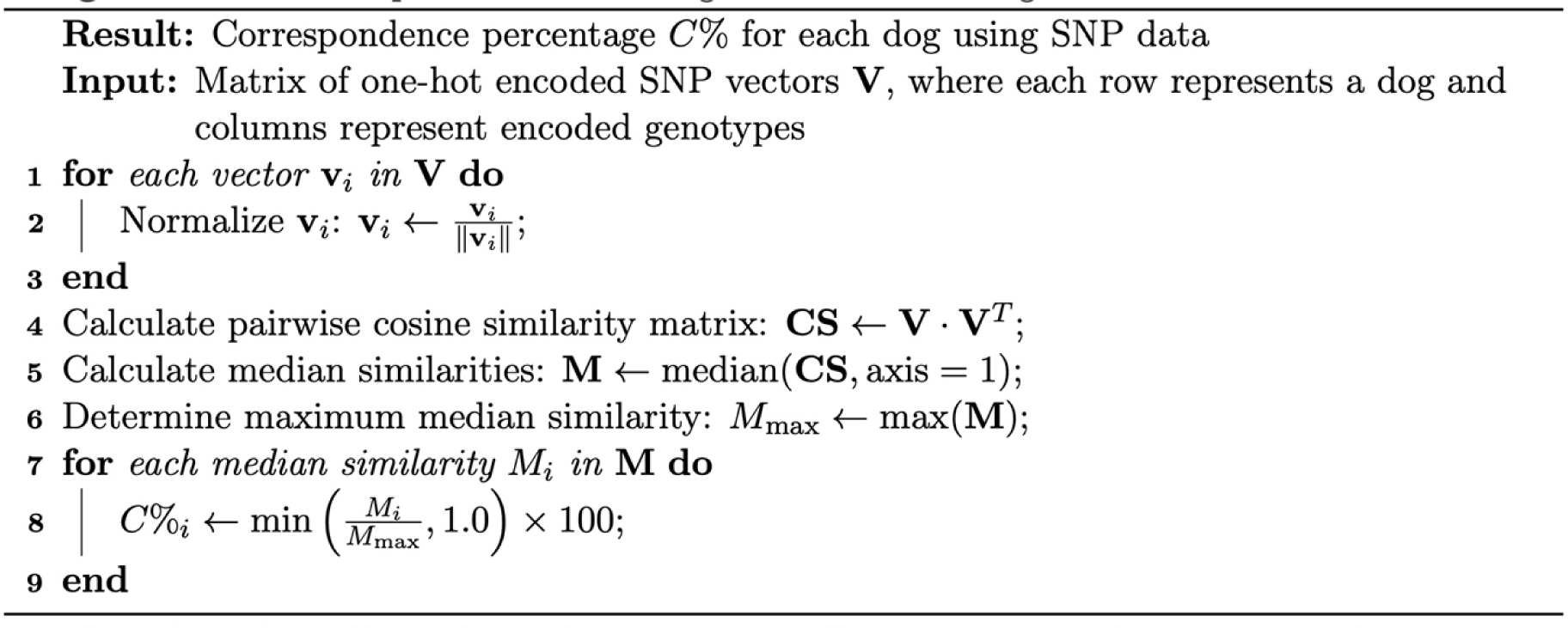

### Expert evaluation

The evaluation of the Kazakh Tazy dog was conducted by an expert with a nationally recognised qualification. The methodology was adapted to evaluate native breeds taking into account their current status and strictly adhered to the breed standard for the Kazakh Tazy approved by the Decree of the Ministry of Ecology and Natural Resources of the Republic of Kazakhstan dated 30 March 2023 No. 101, as well as the guidelines set out in the Rules for Testing Dogs at Dog Shows adopted by the Club of Bloodhound Breeders’ on 29 February 2000. In all cases, the assessment was based on the appearance of the dog in the photos.

In the evaluation of the breed, the following gradations were used: high breed, typical, not typical (with signs of crossbreeding), as well as gradations in the evaluation of conformation: normal (without disadvantages), has a number of disadvantages; has a number of defects and faults; has disqualifications. Taking into account the evaluation of the breed and conformation, a final score was awarded in the following gradations: excellent (5 points), very good (4 points), good (3 points), satisfactory (2 points), low score (1 point). The most important indicator for the final evaluation was the level of the breed, which was primarily determined by the shape of the head.

### Correlation analysis

The Pearson correlation coefficient, which indicates the degree of linear relationship between two methods, was calculated using the following formula:

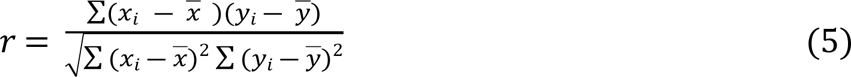

where *x*_*i*_ and *y*_*i*_ are individual data points from two series, and 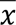 and 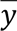 are the means of those series.

The Python package Matplotlib was used to visualise the results [29]. See the pseudocode of the result visualisation in S3 File.

## Results

### Genetic structure

Microsatellite analysis was performed on a sample of 223 Kazakh Tazy dogs for 18 microsatellite loci recommended by the International Society for Animal Genetics (ISAG) for dogs [30]. The allele frequencies at these loci were used to assess genetic variability within the population (Table 1). A total of 195 alleles were detected at the 18 microsatellite loci. All loci exhibited polymorphism, with allele diversity ranging from 6 to 15 alleles per locus. The mean number of alleles per locus (Na) was calculated to be 10.833 ± 0.595, while the mean number of effective alleles per locus (Ne) was determined to be 5.005 ± 0.426. A comparison of the observed heterozygosity (Ho = 0.728) with the expected heterozygosity (He = 0.774) revealed a higher He, which was supported by a positive inbreeding coefficient (F = 0.061).

**Table 1.** Polymorphism of 18 microsatellite markers of the 223 Kazakh Tazy dogs.

|  | Na | Ne | Ho | He | F | ChiSq | P |
| --- | --- | --- | --- | --- | --- | --- | --- |
| AHTk211 | 8 | 3.073 | 0.610 | 0.675 | 0.096 | 28.578 | 0.434 |
| CXX0279 | 12 | 4.349 | 0.749 | 0.770 | 0.028 | 70.359 | 0.334 |
| REN169O18 | 9 | 5.728 | 0.762 | 0.825 | 0.076 | 42.605 | 0.208 |
| INU055 | 9 | 3.770 | 0.655 | 0.735 | 0.109 | 35.016 | 0.515 |
| REN54P11 | 10 | 5.886 | 0.816 | 0.830 | 0.017 | 64.021 | 0.033 |
| INRA21 | 10 | 3.625 | 0.749 | 0.724 | -0.034 | 129.354 | 0.000 |
| AHT137 | 13 | 5.918 | 0.771 | 0.831 | 0.072 | 122.557 | 0.001 |
| REN169D01 | 15 | 8.366 | 0.870 | 0.880 | 0.012 | 83.302 | 0.942 |
| AHTh260 | 14 | 4.276 | 0.717 | 0.766 | 0.063 | 165.859 | 0.000 |
| AHTk253 | 9 | 2.994 | 0.561 | 0.666 | 0.158 | 91.073 | 0.000 |
| INU005 | 14 | 4.919 | 0.762 | 0.797 | 0.043 | 705.658 | 0.000 |
| INU030 | 6 | 3.160 | 0.619 | 0.684 | 0.095 | 30.328 | 0.011 |
| FH2848 | 8 | 5.025 | 0.771 | 0.801 | 0.037 | 58.571 | 0.001 |
| AHT121 | 12 | 8.793 | 0.825 | 0.886 | 0.069 | 91.089 | 0.022 |
| FH2054 | 14 | 5.697 | 0.794 | 0.824 | 0.037 | 489.319 | 0.000 |
| REN162C04 | 10 | 5.035 | 0.780 | 0.801 | 0.026 | 65.038 | 0.027 |
| AHTh171 | 12 | 7.148 | 0.776 | 0.860 | 0.098 | 169.428 | 0.000 |
| REN247M23 | 10 | 2.324 | 0.516 | 0.570 | 0.095 | 149.821 | 0.000 |
| Mean | 10.833 | 5.005 | 0.728 | 0.774 | 0.061 |  |  |
| SE | 0.595 | 0.426 | 0.023 | 0.020 | 0.011 |  |  |

Hardy-Weinberg equilibrium (HWE) estimates showed deviations from equilibrium at 13 of the loci (Table 1). The specific loci that showed significant deviations at P < 0.05 included REN54P11, INU030, AHT121 and REN162C04, while the loci INRA21, AHT137, AHTh260, AHTk253, INU005, FH2848, FH2054, AHTh171 and REN247M23 showed stronger deviations at P < 0.001.

The PCoA revealed three significant axes: axis one explained 4.93% of the existing variation, axis two 4.23% and axis three 3.91% (Fig 1).

**Fig 1.**
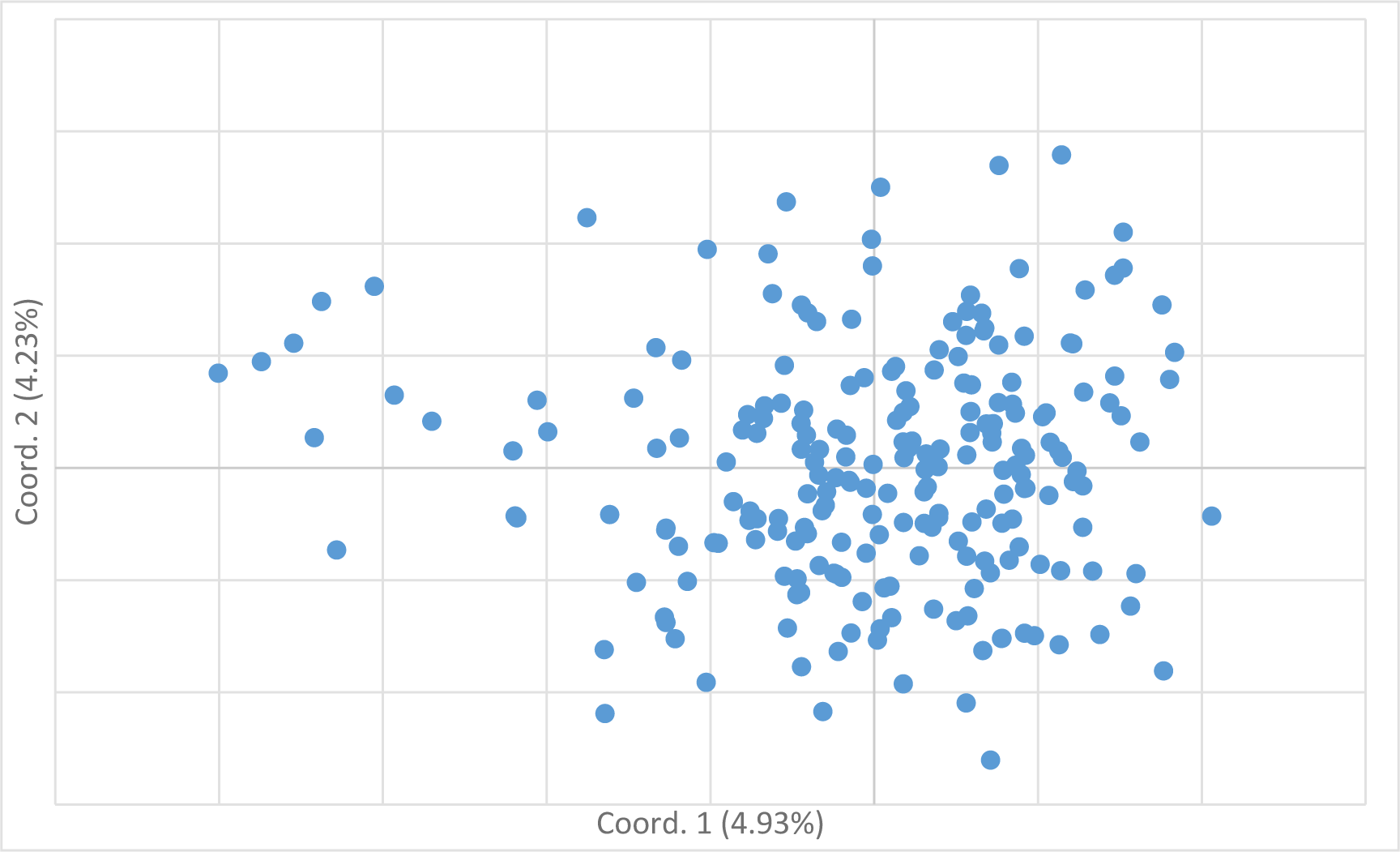
223 Kazakh Tazy dogs clustered on the basis of PCoA using individual microsatellite genotypes.

Based on the average values of the logarithm of the likelihood function and the dispersion of the estimates obtained in ten runs of the program STRUCTURE, the optimal number of clusters was equal to three (K = 3, Fig 2).

**Fig 2.**
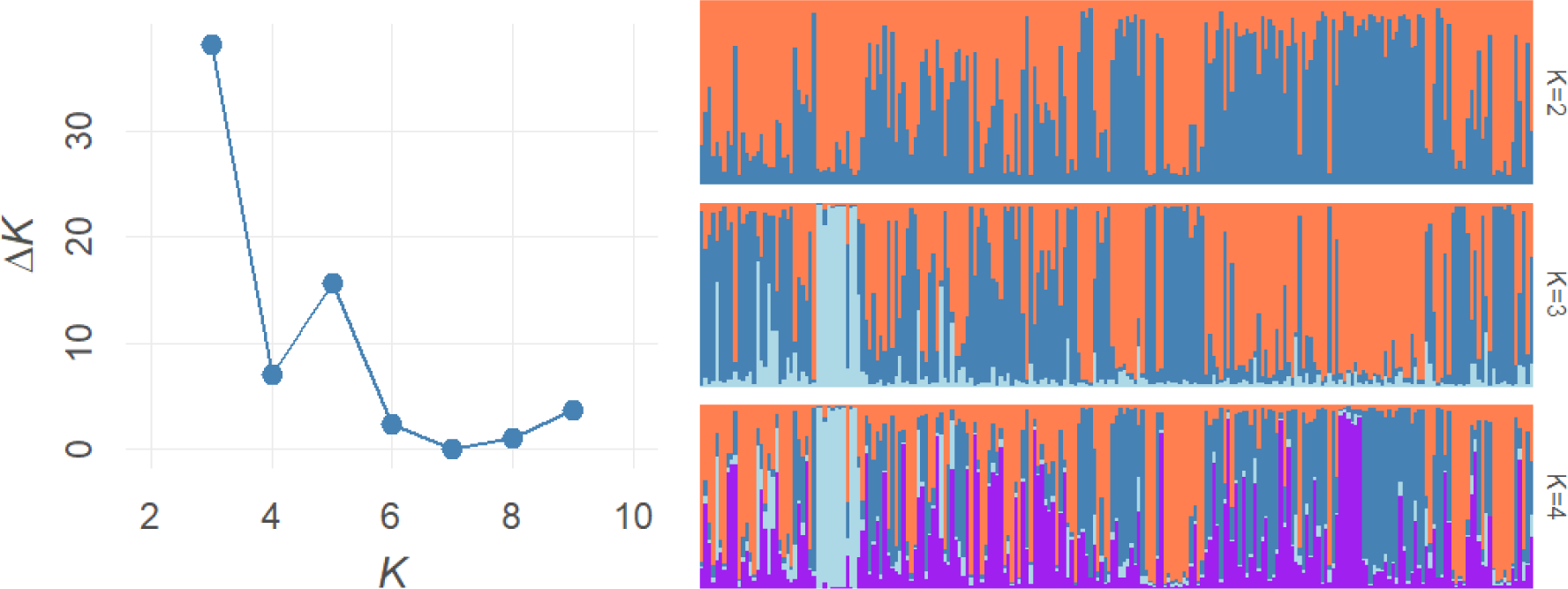
Bayesian clustering on the microsatellite dataset of 223 Kazakh Tazy dogs. the left figure indicates the Delta K results; the right figure indicates - SRTUCTURE plot.

### Results of the expert judgement

A total of 123 dogs out of a total of 223 dogs were evaluated by an expert (S1 Table). Three dogs were assessed as high breed (2.44%), 90 as typical (73.17%) and 30 as not typical (with signs of crossbreeding) (24.39%). Two out of three high breed dogs were classified as having no disadvantages (1.63% of 123 dogs). 118 out of 123 dogs (95.93%) had a number of defects and faults. For three dogs it was not possible to judge the conformation based on photos (2.44%). The average final score was 2.66±1.05. The range and distribution of final score are shown graphically in Figs 3a and 3b. The predominant final score was 3 (71 dogs/57.72%). Only 19 dogs received the high final scores of 4 and 5 (15.45%).

**Fig 3.**
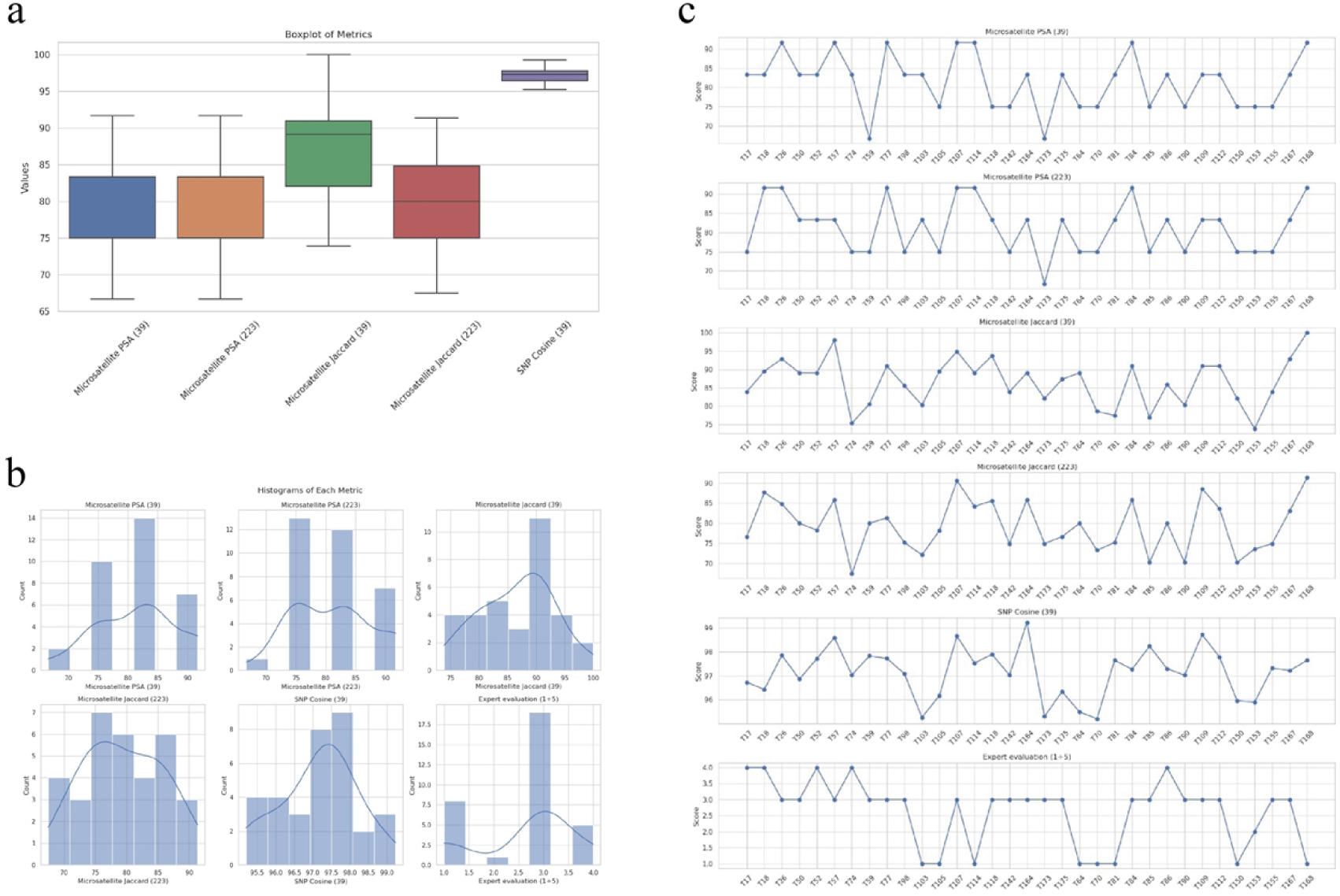
Evaluation of Kazakh Tazy dogs based on the proposed algorithm and expert judgement. (a) - boxplot of the distribution of the breed correspondence value for Kazakh Tazy based on the microsatellite PSA (39) and (232), the microsatellite Jaccard (39) and (232), and the SNP cosine methods; (b) – histograms of the distribution of the breed correspondence value based on the microsatellite PSA (39) and (232), the microsatellite Jaccard (39) and (232), and the SNP cosine methods, and expert judgement; (c) - distribution of the breed corresponding value based on the microsatellite PSA (39) and (232), the microsatellite Jaccard (39) and (232), and the SNP cosine methods and expert judgement for 33 Kazakh Tazy dogs.

### Evaluation of the accuracy of the algorithm

The validation of the proposed algorithm was performed by applying it to microsatellites and SNPs. Since SNP data were available for a cohort of 39 Kazakh Tazy dogs, for fairness of comparison, we calculated the correspondence percentage using the PSA and Jaccard indices based on microsatellite data specifically for two different sample sizes, 39 and 223 Kazakh Tazy dogs. For clarity, the methods were labelled as follows: microsatellite PSA (39), microsatellite PSA (202), microsatellite Jaccard (39), microsatellite Jaccard (202) and SNP cosine methods. The categorical values representing the obtained correspondence percentages can be found in S2 Table. The range and distribution of the obtained values are shown graphically in Figs 3a and 3b, respectively. All microsatellite methods for both sample sizes showed identical variability, which was larger compared to the SNP cosine method. The microsatellite Jaccard (39) method had the highest skewed median, suggesting that this method tends to show high values more frequently.

For 33 Kazakh Tazy dogs with available expert values, SNP and microsatellite data, Fig 3c shows the breed correspondence percentage variability across methods and sample sizes. For the microsatellite PSA method, greater variability in similarity percentages was observed for some dogs with changes in sample size, such as for dog T17, where similarity shifted from 83.33% for a sample of 39 dogs to 75.0% for a sample of 202 dogs, in contrast to the more stable microsatellite Jacard method, where percentages for dog T17 were 83.93% and 76.66% for a sample of 39 and 202 dogs, respectively. Nevertheless, the overall correlation analysis showed agreement between the microsatellite methods at different sample sizes with Pearson correlation coefficients of 0.85 and 0.87 for the PSA and Jaccard methods, respectively (Fig 4).

**Fig 4.**
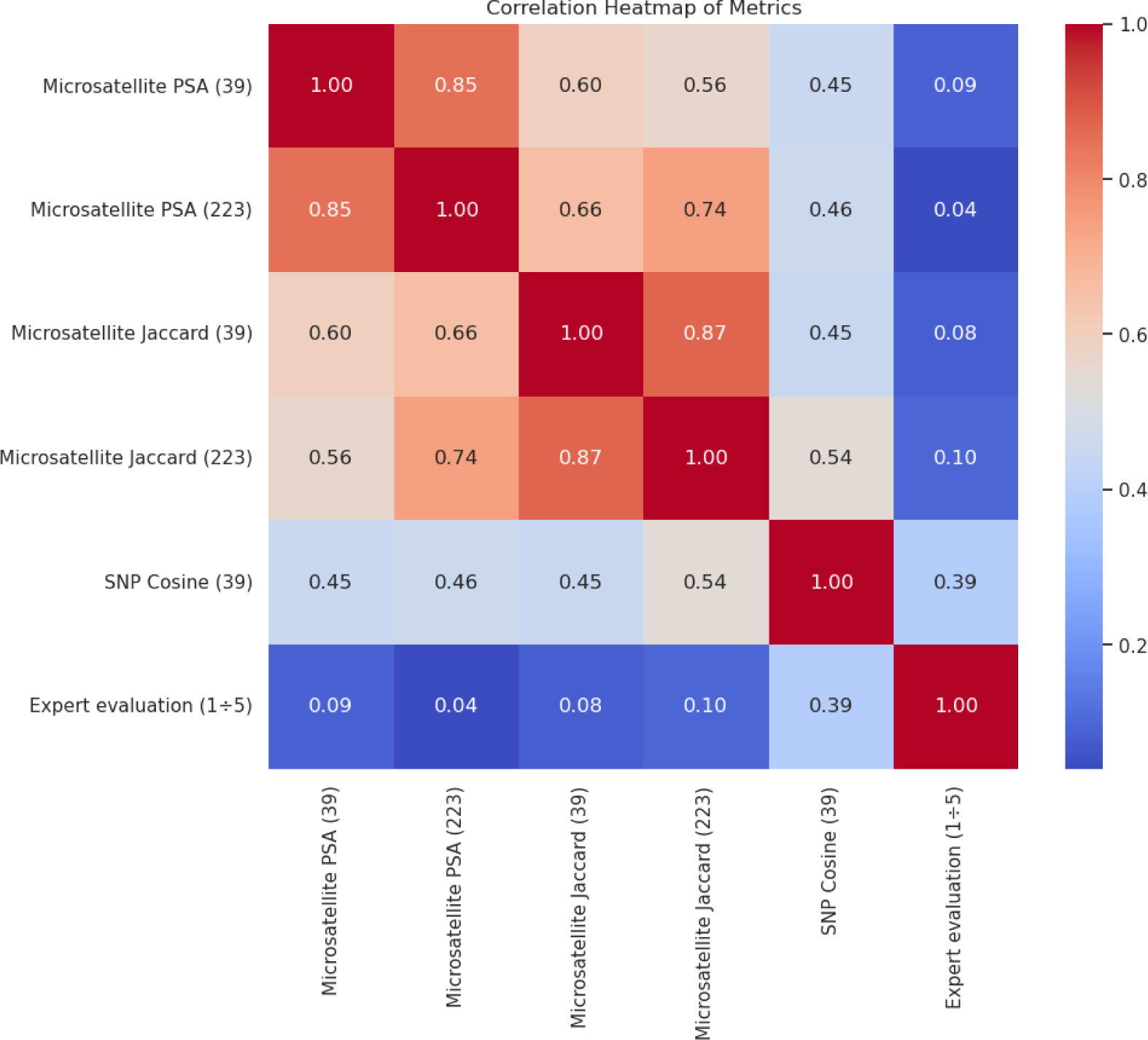
Heatmap of Pearson correlation coefficient matrix. The correlation matrix shows the pairwise correlations between methods and is displayed as Pearson correlation coefficient. The strength of the correlation is colour-coded and ranges from intense blue (no/weak correlation) to intense red (strong correlation).

The correlations between the microsatellite PSA and the Jaccard method for both sample sizes show a moderate degree of linear relationship, which tends to be stronger for the larger sample size: the Pearson correlation coefficients between the microsatellite PSA and the microsatellite Jaccard methods were 0.60 and 0.74 for sample sizes of 39 and 232 dogs, respectively. The other group of moderately correlating methods included all microsatellite methods and the SNP cosine method. For the microsatellite PSA, the correlation indices with the SNP cosine method were almost identical for both sample sizes (0.45 and 0.46 for sample sizes of 39 and 232 dogs, respectively). The correlation of the SNP cosine method with the microsatellite Jaccard method increased slightly with larger sample sizes (0.45 and 0.54 for sample sizes of 39 and 232 dogs, respectively). The strongest positive correlation with the expert judgements was observed with the SNP cosine method (0.39). In contrast, the methods based on microsatellites showed only a weak correlation (from 0.04 to 0.10).

## Discussion

In this study, which included a sample almost twice as large as our previous work [20] we confirmed the high genetic diversity within the Kazakh Tazy breed. The study showed comparable values for different parameters of genetic diversity. In particular, the effective population size (Ne) in the previous study was 4.869 versus 5.005 in the current study. The observed heterozygosity (Ho) was previously 0.748 and now 0.728. The expected heterozygosity (He) showed values of 0.769 and 0.774, and the inbreeding coefficient (F) was previously 0.030, whereas it is 0.061 in the current study. The similarly high level of diversity parameters is characteristic of outbred dogs (Ho = 0.729-0.799) [31] while lower values are reported for other sighthound breeds (Ho = 0.61-0.66) [11,32]. In populations characterized by high genetic diversity, the need to manage and maintain certain breed-specific phenotypic and genotypic traits becomes increasingly urgent.

The origin of this study lies in the need for a scientifically rigorous method to quantify the degree of genetic similarity of certain individuals of a breed compared to the genetic profile of the breed. To achieve this, we developed an algorithm that essentially uses a median-based approach to assess genetic similarity. The decision to use the median rather than the mean is particularly important in genetic studies as it is a robust method that is less prone to bias from outlier data.

Since there is no definitive standard for validating an algorithm, we performed a correlation analysis between genetic similarity measures derived from microsatellite and SNP data, suggesting that the different methods share some commonalities in the assessment of genetic similarity. In addition, expert judgements were integrated to verify the accuracy of the algorithm. The correlation between microsatellite methods across different sample sizes was above 80%, indicating that the median genetic similarity score remains stable regardless of sample size variation. The moderate correlation of 60%-74% between the microsatellite methods using PSA and Jaccard indices suggests that they may capture different aspects of genetic similarity. The PSA index is able to highlight common alleles at the same loci and thus capture a fine-grained aspect of genetic similarity. In contrast, the Jaccard index provides a broader view by measuring the common alleles in relation to the total allele pool, producing a more comprehensive picture of genetic overlap. At the same time the moderate correlation between the microsatellite methods and the SNP-Cosine method, ranging from 45 to 54%, was somehow expected. These methods assess genetic similarities from different angles and capture different elements of genetic variance. In addition, the broader genome coverage and advanced genotyping technologies of SNP data seem to offer advantages over microsatellite methods, as the SNP cosine method provides a closer approximation to the expert values, reaching almost 40%. Compared to microsatellites, SNP arrays cover a larger area of the genome and can therefore capture a broader range of genetic factors that influence phenotype. Besides, SNP genotyping technologies have evolved significantly and offer high accuracy and reproducibility, whereas microsatellites have genotyping errors [33].

Overall, while there is a positive correlation, it is not strong enough to claim that all the proposed methods are perfectly aligned. Essentially, the moderate correlation emphasises the complementary nature of these methods in analysing genetic similarity. The current study was limited to calculating genetic similarities based on microsatellite and genome-wide SNP data. Nevertheless, a feature of the proposed approach is its flexibility in implementation. It is designed to be adaptable to different genetic markers (Restriction Fragment Length Polymorphism (RFLP), Random Amplification of Polymorphic DNA (RAPD), Inter Simple Sequence Repeat (ISSR), etc.) and computational environments and can efficiently process large data sets for different breeds of livestock and non-productive animals. This adaptability ensures that the algorithm can be used in a variety of research and practical contexts.

Another strength of the proposed algorithm is that it was developed specifically for individual-centred analysis. The correspondence percentage is an easily understandable and practical measure for breed evaluation. It differs significantly from algorithms that focus on broader population structure analyses and provide less interpretable results for specific breed representatives.

For example, PCA is a widely used technique for understanding the genetic structure of populations and visualising the position of an individual in relation to a defined breed cluster in genetic space [34–37]. It projects individuals into a space that captures most of the variation, and samples that are genetically similar are close to each other in the projected space. Thanks to this property, PCA allows a heuristic assignment of individuals to their main parental population [38] and is closely related to admixture analysis, a powerful tool in genetic studies to determine the proportion of an individual genome that originates from different ancestral populations. In the admixture analysis, summarised statistics on allele sharing, the so-called f-statistics, are calculated [39]. It allows the researcher to classify individuals of unknown ancestry to discrete populations [40,41], can reveal the composite nature of an individual ancestry and is particularly informative in breeds with a history of crossbreeding [42–44]. It is excellent for unravelling the ancestry of individuals and populations [45–48], making it an invaluable tool in studies where understanding the historical composition and migration patterns of populations is crucial. Although the application of these methods in calculating the breed correspondence value for an individual is associated with challenges related to the conversion of visual proximity into a quantifiable metric, two SNP-based methods have recently been presented for the assignment of new individuals to the reference population of the Hungarian Vizsla shorthaired dog breed [44]. Specifically, the PCA distance method calculates the standardised distances of individuals to the median position of breed members defined by the coordinates of the PCA. The Identity-by-State (IBS)-central method calculates the standardised distances based on the identity values, where the reference point is an animal that is genetically closest to all members of the reference population. The implementation of these methods, which use median values similar to our approach, encounters particular challenges when applied to the Kazakh Tazy breed. The very limited pool of dogs with high expert values and well-documented pedigrees going back to the fourth generation, in which all ancestors are proven to be Kazakh Tazy, poses a challenge in creating a robust reference group for genetic comparisons.

The algorithm proposed in this study may thus prove useful for genetic improvement programmes for breeds with incomplete or missing pedigree data, when the effectiveness of other methods of assessing individual breeding value may be limited for this reason. For example, a single-step Genomic BLUP (ssGBLUP) method integrates all simultaneously available phenotypic, pedigree and genomic information to provide genomic performance values for genotyped and non-genotyped individuals via the combined matrix H [49–51]. The matrix H can thus be understood as a modification of regular pedigree relationships by including genomic relationships. This enables the utilisation of all available information in a selection. The ssGBLUP has already been successfully used for several large-scale analyses, including dairy cattle, pigs and chickens [52–57]. The potential of genomic selection to improve behavioural traits in the Labrador Retriever breed and the genomic prediction of orthopaedic traits in various pure and mixed breeds was evaluated using the ssGBLUP method [58,59]. However, in situations with missing pedigree data, this method may encounter challenges such as biased predictions and incompatibility between genomic and pedigree relationship matrices [60].

These problems can be solved with the unknown group (UPG) model, which adjusts the genetic relatedness matrix to include unknown parental groups. It helps in estimating breeding values even when complete pedigree information is not available [60,61]. However, it has recently been reported that the UPG model in ssBGLUP also generates bias because it ignores covariance between genetic groups [62,63]. The effectiveness of the UPG model may also be limited in scenarios with high diversity breeds due to crossbreeding, as the mixed genetic background may not match well with the assumptions of the model.

The Metafounder (MF) model is an extension of the UPG model [60,64] and generates more accurate breeding values [62,63]. It creates ‘metafounders’ that represent the genetic basis of the ancestors of animal groups with unknown pedigrees. The metafounders are used to create a more comprehensive genetic relationship matrix, which is then integrated into genomic analyses such as ssGBLUP. However, the MF model requires accurately representing the ancestral genetic base of the breed. The high degree of genetic diversity of a breed such as Kazakh Tazy can make it difficult to determine the meta-founders and subsequently create a comprehensive genetic relationship matrix, which affects the accuracy of genomic evaluations using the MF model.

The same limitations can occur when applying the algorithm for estimating genetic similarity matrices based on phylogenies [65]. The algorithm provides simulations suggesting that genetic similarity matrices calculated from trees match those calculated from genotypes and provides a unique opportunity when quantitative analyses of genetic traits and analyses of heritability and coherence can be performed directly using genetic similarity matrices in the absence of genotype data or under uncertainty in the phylogenetic tree. However, this approach is limited to the calculation of genetic similarities between species or samples without recombination. Previously, we identified three genetic clusters within the Kazakh Tazy breed [20], which were confirmed in this study with a larger sample and could hypothetically be a consequence of recombination. Thus, it is possible that this algorithm may not fully capture the genetic complexity of such a breed affected by crossbreeding.

Classifiers and machine learning models can be trained to classify individuals into breeds based on their genotypes. Classification confidence or probability can be interpreted as a correspondence percentage [66–68]. Various machine learning methods have been used for breed identification, including Bayesian, Naive Bayes, Support Vector Machine (SVM), k-Nearest Neighbour, Random Forest (RF), Artificial Neural Networks (ANN) and Decision Tree [66–69]. These methods have shown high accuracy in breed identification, with accuracies of over 95% in different scenarios [66,69,70]. Trait selection methods, such as Deep Neural Networks (DNN), Garson and Olden, have been used to identify informative SNP markers for breed assignment [67]. In addition, the use of genomic data and machine learning techniques, such as eXtreme Gradient Boosting (XGBoost) and Stacking Ensemble Learning, has improved the accuracy of breed identification [69].

However, machine learning models for breed classification in genetics face several challenges. Their effectiveness depends on access to large, high-quality data sets. Insufficient or biassed data can lead to inaccurate classifications. These models are also prone to overfitting, especially if the training data is not diverse enough, which can lead to poor performance on new data. In addition, such models, especially those trained only on purebred data, may not accurately classify mixed breeds or those that do not fit well with established breed categories. The process of selecting the most informative genetic markers for classifying breeds is another complex aspect, as the effectiveness of trait selection methods can vary. Furthermore, to maintain the accuracy and relevance of these models, they need to be regularly retrained and updated as new genetic data becomes available, which can require significant resources.

The limitation of the algorithm we developed is the moderate correlation with expert judgements, which indicates areas for improvement. Further work should be carried out to identify breed-specific SNP markers, the use of which could optimise the algorithm. In addition, the algorithm is currently limited to analysing genetic similarities based on microsatellite and SNP data, suggesting that it needs to be extended to include other genetic markers to improve its applicability and effectiveness in different animal breeds.

## Conclusion

To summarise, the proposed approach is perhaps the only one that is possible in breeding scenarios without a reference population, missing pedigree, undefined meta-founders and high genetic diversity. It can be used in breeding programmes aimed at achieving or maintaining specific breed standards, where the clear and understandable median value of similarity can serve as a genetic benchmark for breed standardisation. Initial comparisons with two types of genetic markers and expert judgement indicate promising improvements in accuracy. Future efforts will focus on extending the dataset to different breeds to improve the applicability of the algorithm and address the current limitations.

## Acknowledgements

We would like to thank the dog breeders, owners and Public Foundation «Tailed Paradise» who provided us with samples and information about this unique breed. Our special thanks to the veterinarians and members of “Kansonar” who helped with sampling.

## Supporting Information

**S1 Table. Expert evaluation of the Tazy dogs.**

**S2 Table. Breed correspondence percentages for 39 Tazy dogs.**

**S3 Table. Microsatellite dataset for Tazy dogs.**

**S4 Table. SNP dataset for Tazy dogs.**

